# Hybrid multisite silicon neural probe with integrated flexible connector for interchangeable packaging

**DOI:** 10.1101/2021.02.02.429112

**Authors:** Ashley Novais, Carlos Calaza, José Fernandes, Helder Fonseca, João Gaspar, Luis Jacinto

## Abstract

Multisite neural probes are a fundamental tool to study brain function. Hybrid silicon/polymer neural probes, in particular, allow the integration of complex probe geometries, such as multi-shank designs, with flexible biocompatible cabling. Despite these advantages and benefiting from the highly reproducible fabrication methods on both silicon and polymer substrates, they have not been widely available. This paper presents the development, fabrication, characterization, and in vivo electrophysiological assessment of a hybrid multisite multi-shank silicon probe with a monolithically integrated polyimide flexible interconnect cable. The fabrication process was optimized on wafer level and several neural probes with 64 gold electrode sites equally distributed along 8 shanks with an integrated 8 μm-thick highly flexible polyimide interconnect cable were produce. To avoid the time-consuming bonding of the probe to definitive packaging, the flexible cable was designed to terminate in a connector pad that can mate with commercial zero-insertion force (ZIF) connectors for electronics interfacing. This allows great experimental flexibility since interchangeable packaging can be used according to experimental demands. High-density distributed in vivo electrophysiological recordings were obtained from the hybrid neural probes with low intrinsic noise and high signal-to-noise ratio (SNR).

## 1. Introduction

Advances in microengineering and electromechanical systems (MEMS) technology and microfabrication methods and materials have enabled the development of integrated high-density silicon (Si)-based neural probes for neuroscience applications [1]–[8]. The superior adaptability and reproducibility of Si microfabrication processes has allowed the continuous refinement of probes’ geometry parameters which in turn has led to improved surgical implantation procedures, increased mechanical stability and higher signal to noise ratio (SNR) neural recordings. Hence, it is currently possible to monitor the simultaneous activity of dozens to hundreds of individual neurons in multiple sites with these probes, which has been contributing to our understanding of information processing and coding in the brain and the development of more effective brain-machine interfaces [9]–[12].

More recently, there has been a rising interest in flexible neural probes and interconnect cables since they can reduce the mechanical mismatch between probe and brain tissue, permit fully implanted biocompatible cabling and facilitate integration with flexible/organic electronics [13]–[16]. However, and despite these advantages, flexible probes can buckle during brain insertion and require cumbersome mechanical rigidity augmentation strategies to increase the probe’s buckling force threshold [14], [17], [18] during insertion/implantation. Consequently, while probes fabricated on compliant substrates such as polyimide (PI), parylene C or SU-8 can be seamlessly integrated with flexible interconnect cabling they typically have limited geometry options. In particular, polymer probes with multiple-shank designs require complex augmentation strategies and have rarely been pursued [17], [19]–[22]. Additionally, the polymer layers of integrated all-polymer probes are usually thicker than the ones used for cabling only since they must structurally support the implantable portion of the probe, thus leading to thicker and less flexible cabling.

Hybrid silicon/polymer neural probes that combine high-definition silicon probes with monolithically integrated thin polymer flexible interconnect cables can be an alternative solution, but have received limited attention [23]–[27]. This approach takes advantage of microfabrication processes on both silicon and polymer substrates and allows the fabrication of small-footprint silicon probes, including multisite/multi-shank designs that do not require additional brain insertion aids, while retaining thin fully flexible cabling for electronics interfacing.

Nevertheless, even when flexible cabling is integrated, either in silicon or polymer substrates, there is still the need to perform a time-consuming packaging step of wire- or flip-chip bonding of the cable interconnect pads to a PCB used for electronics interfacing. Considering that different experimental demands and applications may require different probes, there is also the need for specific PCBs and electronics interfaces for each different probe. The inexistence of a standard for probe interfacing, while allowing some flexibility, also means that labs must invest in different types of connectors and packaging options for each probe and experiment, typically dictated by non-experimental requirements. Although fabrication techniques have been evolving towards more cost effective high-yield approaches, the use of Si probes is still forbidding for many neuroscience labs, especially for chronic experiments, with the cost of different packaging options adding up. This has limited the use and dissemination of probes, especially of those developed in labs [28].

To facilitate wider dissemination and use, we propose here a new fabrication process for hybrid multisite silicon probes with integrated polyimide flexible interconnect cabling that allows inter-changeable packaging options. By combining optimized fabrication processes for Si and PI, we fabricated several small-footprint multi-shank probes with a higher channel count (64 electrode sites) and thinner flexible polymer interconnect cabling (8 μm thick) than previously reported hybrid silicon/polymer probes [23]–[27], [29]. And by designing an integrated open-ended flexible connector pad that can mate with commercial zero-insertion force (ZIF) connectors on the PCB side, definitive packaging for these probes is avoided allowing great experimental flexibility. With this design, the same probe can be easily connected to different custom-designed PCBs for each required application, or different probes, with different geometries and layouts, can use the same interface PCB. Additionally, by avoiding definitive bonding of the probe to the connector, the speed of fabrication increases while overall costs decrease. This also leads to improved wafer utilization which, when combined with optical lithography near its patterning limits, as we show here, can increase the number of probes fabricated per wafer, further lowering costs. Figure 1 shows a schematic of our hybrid multisite silicon neural probe with integrated flexible polyimide cable. This paper describes the design, fabrication and *in vivo* electrophysiological assessment of these probes.

**Figure 1:**
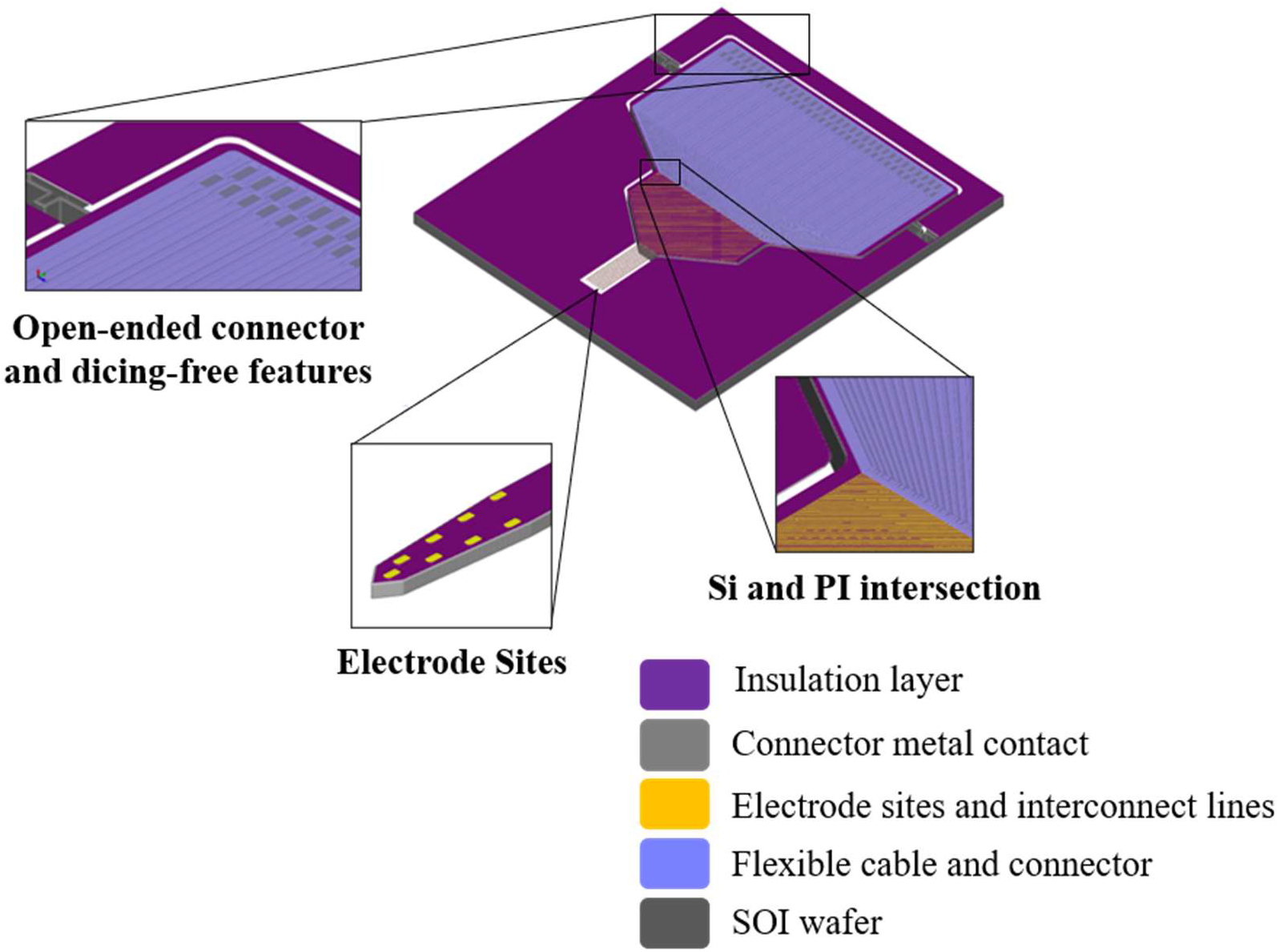
3D schematic of the neural probe.

## 2. Material and Methods

### 2.1 Microfabrication of Neural Probe

The neural probe was fabricated using standard semiconductor micromachining processes. Briefly, 200mm SOI (silicon-on-insulator) wafers (15 μm device layer, 2 μm buried oxide, 625 μm handle wafer) (SVM, Santa Clara, USA) were used as substrate (Figure 2a). Back-side was protected with one extra micron of SiO_2_ deposited by plasma-enhanced chemical vapor deposition (PECVD) as hard mask, and a layer of 500 nm Al_2_O_3_ was sputtered for front-side passivation. A metal stack layer of 15nm TiW/150nm Au/5nm Cr was then sputtered and patterned via reactive ion etching (RIE) (Figure 2b). This was followed by deposition of a 500nm layer of Al_2_O_3_ for passivation defined by wet etch, and back-side was patterned for DRIE (Figure 2c). To proceed with the definition of the polyimide (PI) connector, electrode sites were first protected with a stack of 500 nm AlSiCu defined via wet etch. A 500 nm SiO_2_ sacrificial layer for PI release was then patterned via RIE, followed by a 3.75 μm-thick layer of PI (PI-2611, HD MicroSystems, New Jersey, USA) which was spin-coated and cured at 250° C for 14 hours and etched via RIE to open vias (Figure 2d). Then, the AlSiCu metal stack (1000 nm AlSiCu/150nm TiW/200nm AlSiCu/50nm TiW) of the inter-connector was patterned via metal RIE followed by deposition of another 3.75 μm-thick layer of PI and connection pads were patterned via PI RIE for connector definition (Figure 2e). To define the probe shanks the Al_2_O_3_ passivation layer was etched to allow Si DRIE on the front-side. Al was, then, wet-etched to remove protection of device area and Cr was wet-etched to expose the Au sites (Figure 2f). After 15 μm probe definition, the front-side was protected using Cool Grease (AI Technology Inc., New Jersey, USA) for wafer bonding to a handling wafer. The 625 μm of Si on the back-side was etched via DRIE (Figure 2g), followed by hydrogen fluoride (HF) vapor etch to remove the buried oxide layer and release the PI connector (Figure 2h). Finally, handling wafer was released in water bath (60 °C) and cleaned with acetone. Since dicing the wafer reduces wafer yield, a dicing-free process [30] was implemented for individual probe release from the wafer. For that purpose, a 50 μm-wide trench around the device was patterned with DRIE on front- and back-side. U-shaped breakout beams connect the side of the probes to the bulk wafer allowing safe wafer handling during process and effortless device release by simply breaking the beams with tweezers.

**Figure 2.**
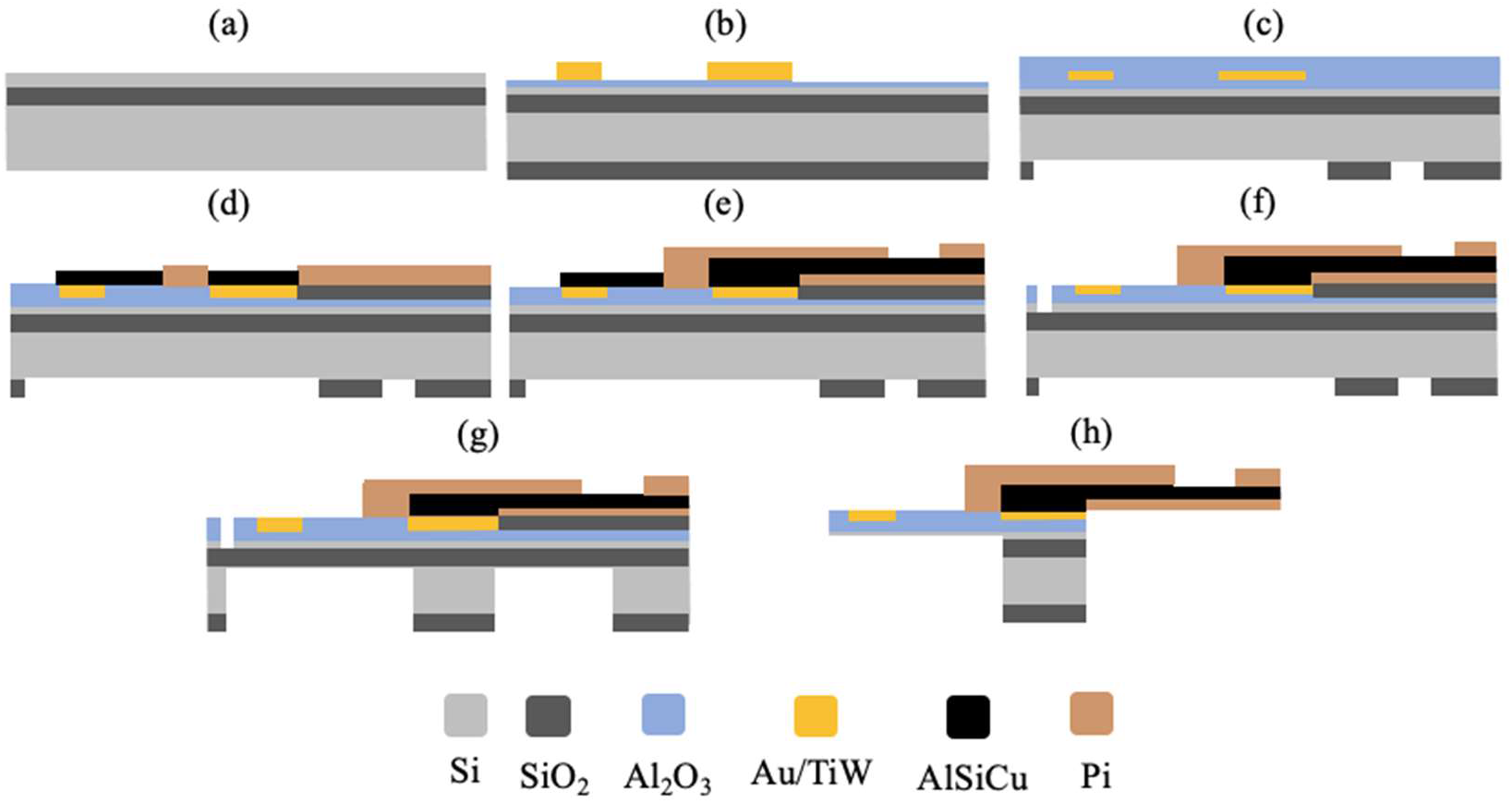
Neural probe main fabrication steps. a) SOI Wafer; b) Au/TiW thin-film patterning and back-side SiO_2_ layer deposition; c) Font-side Al_2_O_3_ passivation and back-side SiO_2_ layer patterning; d) electrode sites protection and SiO_2_ and PI patterning; e) Interconnector AlSiCu and PI patterning; f) Front-side Si DRIE, g) Back-side Si DRIE and h) HF SiO_2_ release. NB: not to scale.

The fabrication process was also optimized at the wafer level, which allows the scale-up of the fabrication of probes with different geometries and layouts within the same wafer.

### 2.2 Neural Probe Packaging

For signal acquisition for the impedance measurements and *in vivo* electrophysiological recordings reported here the probe was packaged with a custom-designed PCB with a FPC ZIF connector (FH39A-67S-0.3SHW, Hirose Electric) on the probe side and two omnetics connectors (A79022-001, Omnetics Connector Corporation) on the acquisition system side. The open-ended polyimide connector pad of the integrated flexible cable was inserted into and secured to the ZIF connector for signal acquisition.

### 2.3 Electrical Impedance Analysis

Prior to electrode site impedance measurements, the tips of the probe shanks were immersed in a solution of 50 mM potassium hydroxide (KOH) and 25% hydrogen peroxide (H_2_O_2_) for 10min, as described in [31] to remove fabrication process residues and obtain clean gold electrode sites. Shanks were then rinsed in abundant milli-Q water. Electrode site impedance was measured with nanoZ (White Matter LLC) in a phosphate buffered saline solution (PBS 1x) at 1kHz.

### 2.4 In Vivo Electrophysiological Recordings

All animal procedures complied with the European Union Directive 2016/63/EU and the Portuguese regulations and laws on the protection of animals used for scientific purposes (DL No 113/2013). This study was approved by the Ethics Subcommittee for the Life Sciences and Health of University of Minho (ID: SECVS 01/18) and the Portuguese National Authority for Animal Health (ID: DGAV 8519).

Wild-type mice were anesthetized by intraperitoneal injection mix of ketamine (75mg/Kg) and medetomidine (1mg/Kg). Surgical procedure consisted of exposing the skull following a midline skin incision and drilling a burr hole above the motor cortex (at 1.0 mm AP and 1.5 mm ML from bregma [32]). Following dura removal, a 5 cm long, 8-shanks neural probe with 64 electrode sites connected to the custom-designed PCB via the FPC ZIF connector was attached to a micrometric stereotaxic arm (1760, Kopf Instruments). The PCB was then connected to an headstage (RHD2132, Intan) for signal acquisition. Probe was lowered into the brain, through the burr hole, to a depth of −0.5 mm (DV) from the brain surface. A stainless-steel ground screw, positioned in another burr hole at the back of the skull, was connected to the PCB ground pad. Spontaneous extracellular neuronal activity signals were acquired with an Open Ephys acquisition system at 30 Ks/s [33].

Extracellular neuronal recordings were analyzed with custom-written Matlab code (Mathworks) and initial spike sorting was performed using JRClust [34]. Recordings were filtered between 0.6 and 6 kHz and spikes were detected using an amplitude threshold at least 5 times higher than background noise standard deviation. Manual curation of single unit clusters, after JRClust initial automatic spike sorting, was performed by visual inspection of inter-spike interval histograms, auto- and cross-correlograms and clusters’ spike waveforms.

Signal-to-noise ratio (SNR) was calculated using the formula: 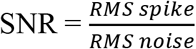, where RMS spike is the average of the root mean square of 1ms windows centered at the peak of each detected spike and RMS noise is the average standard deviation of the portions of the extracellular signal where spikes did not occur.

## 3. Results

### 3.1 Design and Fabrication of Neural Probe

Figure 3a shows an example of a fabricated neural probe. The probes were designed and fabricated with 8 shanks with a pitch of 200 μm and 8 electrode sites per shank, for a total of 64 electrode sites. Each shank has a maximum width of 54 μm and a thickness of 15 μm and three different shank lengths were produced (2.5, 5 and 10 mm, Figure 3b). Gold electrode sites on shank tips have an area of 72 μm^2^ (6 μm × 12 μm) and are distributed vertically in two columns along the two edges of each shank with a vertical pitch of 20 μm (Figure 3c).

**Figure 3.**
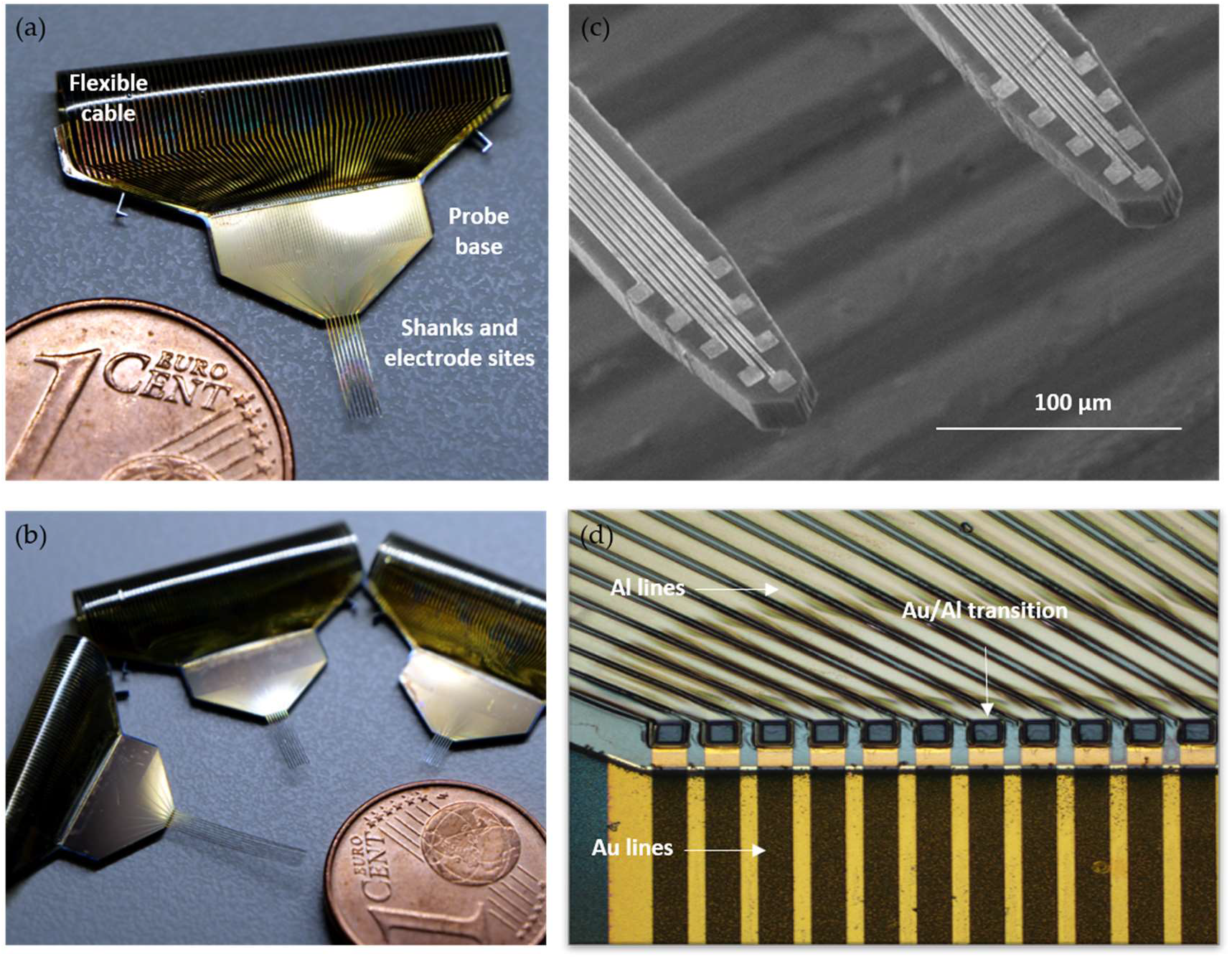
Fabricated neural probe: (**a**) Photograph of a 5 mm long silicon probe with polyimide flexible cable; (**b**) Probes with three different shank lengths, 2.5, 5 and 10 mm; (**c**) SEM image of two shanks of a neural probe, with 8 gold electrode sites and respective interconnect lines in each shank; (**d**) Detail of the intersection zone of gold (Au) and aluminum alloy (Al) interconnect lines from the silicon and polyimide portions of the probe, respectively.

The metal interconnect lines arising from each electrode site travel up along the shanks and end in larger gold metal pads (100 × 100 μm) on the base of the Si probe. These pads form the transition zone where the electrodes’ gold interconnect lines contact with the flexible polyimide (PI) cable aluminum alloy intermetal lines (Figure 3d). The gold interconnect metal lines have a width of 2 μm which equals the patterning resolution limit of the low-cost lithography process implemented here. At 54 μm, shanks have the minimum possible width to accommodate all electrode sites and the interconnect metal lines. Energy dispersive X-ray spectroscopy (EDX) structural analysis of the electrode sites and interconnect metal lines can be found in Appendix A (Figure A1).

The highly flexible 8 μm-thick integrated polyimide interconnect cable is 3 cm long but could be extended to any desired length. The interconnect cable ends in a pad array with a custom format for effortless insertion into a commercial ZIF connector. By not require definitive bonding to a PCB, the open-ended connector allows interchangeable packaging according to experimental needs.

### 3.2 Electrical Impedance of Neural Probe

Following connection of the probe to the custom PCB and electrode site gold surface cleaning, electrode electrical impedance was measured. The average electrode impedance measured at 1kHz in phosphate-buffered saline (PBS) was 441.97 ± 47.98 kΩ.

### 3.3 In vivo Electrophysiology

To assess the fabricated probes’ performance in the context of an *in vivo* experiment, brain electrophysiological extracellular recordings were performed, as depicted in Figure 4a. Neuronal activity from the motor cortex (M1) of anesthetized mice was recorded from probes with high signal-to-noise ratio (SNR) across electrode sites and permitted subsequent reliable spike sorting. Figure 4b shows example traces of neuronal activity recorded from eight electrode sites in one shank of the probe. The mean RMS of the recorded extracellular signals across electrode sites was 5.29 ± 0.6 μV with an SNR_Voltage_ of approximately 7.1 ± 0.6. Due to the excellent SNR, it was possible to detect high-amplitude spikes (Figure 4d) which were then sorted into single unit clusters. Figure 4e shows the waveforms of 4 single units isolated from one shank of the probe during a 5-minute recording.

**Figure 4.**
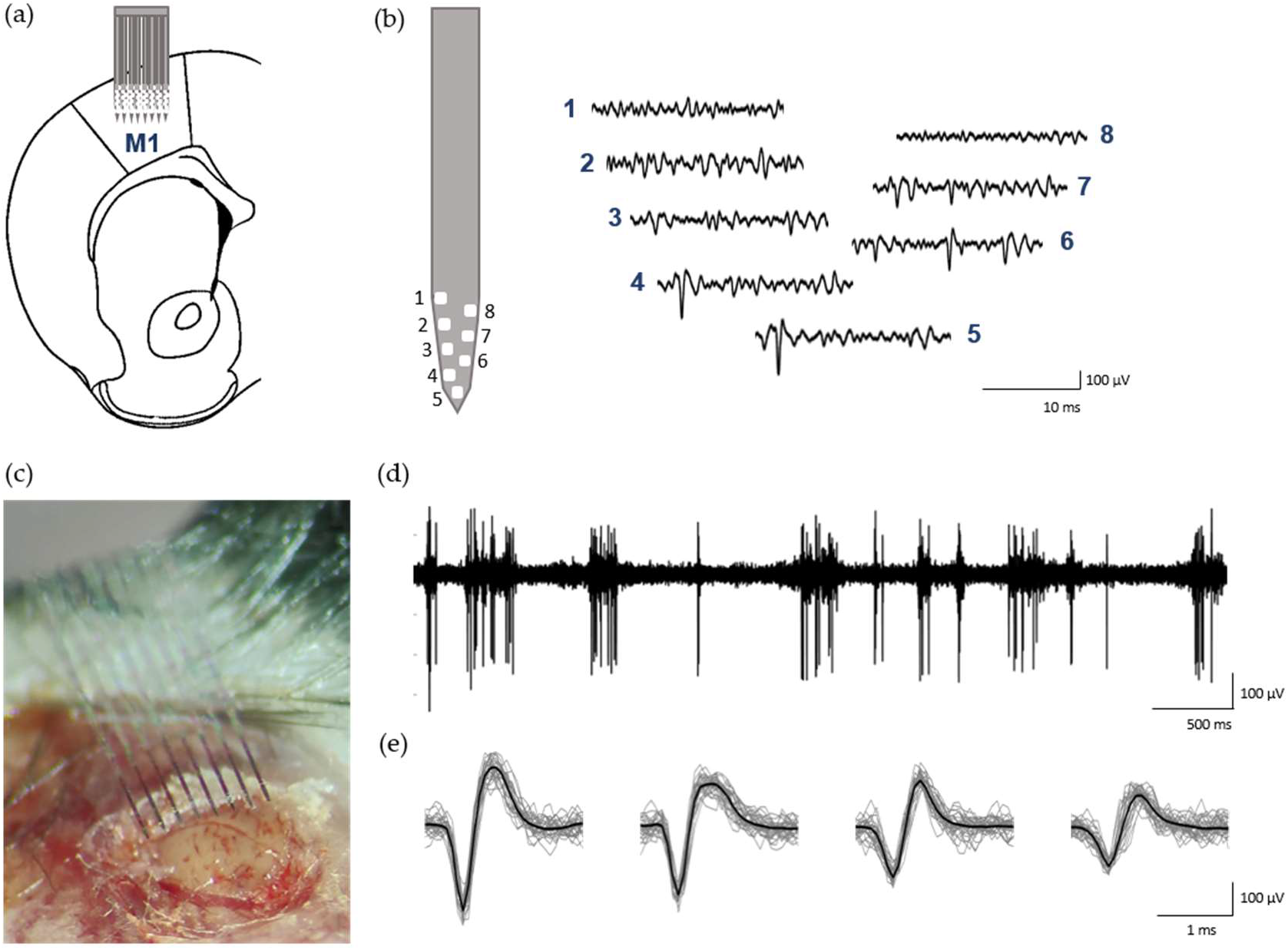
*In vivo* recordings in the mouse primary motor cortex (M1): (**a**) Schematic representation of probe insertion in M1; (**b**) Example of neuronal activity simultaneously recorded from 8 electrode sites from shank 1 of the probe (signals band-pass filtered between 0.3 and 6 kHz); (**c**) Surgical procedure for neural probe insertion in M1; (**d**) Example signal trace of neuronal activity recorded from one electrode site with high SNR action potentials (filtered 0.3 – 6 kHz); (**e**) Isolated single-units from the same recording, showing mean waveform (black) and the first 100 spike waveforms (grey);

## 4. Discussion

A hybrid silicon neural probe with integrated polyimide flexible cabling and open-ended connector for interchangeable packaging is presented. The monolithic fabrication process described permitted the integration of a multi-shank multisite 64-channel silicon (Si) neural probe with an 8 μm-thick highly flexible polyimide (PI) cable. When compared with previously published hybrid neural probes with flexible cables [23]–[26], [29], the one described here presents a significantly higher number of electrode sites and a thinner interconnect cable. Additionally, an open-ended connector pad was designed at the end of the flexible cable to permit the use of any desired printed circuit board (PCB) for probe interfacing, as long as a ZIF connector is present on the PCB side. This allows great experimental flexibility since the packaging can be easily changed to meet experimental demands without any necessary alterations to the probe or cable connector.

The employed monolithic fabrication process relies on optimized fabrication processes for Si and PI can be divided in 2 parts: fabrication of the Si probe and fabrication of the integrated flexible PI interconnect cable. The substrate of choice for the neural probe was a SOI (silicon-on-insulator) which allowed precise control of the neural probes’ implantable shank thickness and that other parts of the probe remained at full wafer thickness [35]. The thickness of the shanks was determined by the SOI device silicon layer, which was 15 μm, while the probe’s base remained at wafer thickness (approx. 640 μm) to provide support for safer handling. Other hybrid probes with integrated flexible interconnect cables that did not use SOI wafers reported considerably higher probe thicknesses (50-150 μm) [23], [24] which can cause increased brain tissue damage and reactive responses upon insertion/implantation [36], [37]. At 15 μm thick, the shanks on our probes are small enough to minimize tissue displacement and brain damage upon insertion and sufficiently rigid to avoid buckling and to provide great mechanical stability.

Alumina (Al_2_O_3_) was chosen for neural probe passivation. Although SiO_2_ is most commonly used for this purpose, it was not compatible with the implemented fabrication process since it was used as the sacrificial layer for PI release. Additionally, alumina is biocompatible and chemically stable for chronic implantation [38]. Parylene or PI have also been previously used as insulators for other Si neural probes with flexible integrated interconnect cables [24], [26], but it significantly increased the thickness of the implanted portion of those probes when compared with ours. Although using PI as a passivation layer could facilitate the integration of our PI cable, the 500 nm thick alumina passivation layer used ensured that probe thickness was kept to the minimum (which would have otherwise increased if a 3-5 μm thick polyimide passivation layer was used). The undesirable increase in thickness would also defeat the purpose of using a SOI wafer for precise control of reduced shank thickness.

Each shank on the neural probe has 8 gold electrode sites distributed along the two edges of the shank. This electrode site layout increases spatial sampling while also permitting over-representation of neural activity across different nearby sites, which facilitates spike sorting and increases single-unit yield and separation [7], [39]. Sputtered gold was the chosen metal for the electrode sites, but the same fabrication process could be employed using another metal with suitable electrical impedances. Gold sites have the advantages of being biocompatible and not requiring additional electrodeposition procedures to lower impedances to the desired range, at least with the site dimensions used here. The measured mean electrical impedance of the electrode sites was within the optimal range for neural recordings with high SNR [40] and the recorded neural signal RMS was comparable to other state-of-the-art neural probes [3], [4]. In the *in vivo* electrophysiological assessment of the neural probe, several single units with high SNR were reliable isolated from all shanks.

To avoid time-consuming post-fabrication bonding processes to connect a flexible interconnect cable to the silicon probe, as for ex. in [41], a microfabrication process based on PI was used here to monolithically integrate a polyimide interconnect cable in the probe. PI was the chosen substrate because of its conformational rigidity, dielectric properties and process compatibility [42]. An aluminum alloy (AlSiCu) was used as interconnector metal on the PI cable because it is not only more affordable than a noble metal but also displays good adhesion properties to Si and PI and low residual stress [43]. The area of the interconnect metal pads of the Si/PI transition zone, both on the probe and cable sides, as well as the width of interconnect lines on the probe base, could be further reduced to create a smaller probe base which would be more amenable to chronic brain implantation. Nevertheless, the 8 μm thick polyimide cable is the thinnest ever reported for a hybrid silicon/polymer neural probe which would be advantageous for cabling bio-implantation.

The flexible interconnect cable was also designed to terminate in a pad array that can be easily connected to any PCB for electronics interfacing with a ZIF connector. The connector package creates an interface between the electrode sites and the external electronics for signal acquisition. Si probes typically have rigid interconnects that require time-consuming wire- or flip-bonding processes to physically bond them to an interfacing PCB. Additionally, commercial probes typically also have a limited number of packaging options an experimenter can choose from. The approach presented here avoids definitive physical bonding of the probe to the electronics interface and allows interchangeable packaging according to the experimental/application demands. With this approach, the same neural probe can be easily connected to different custom-designed PCBs for each required application, or different probes, with different geometries and layouts, can use the same interface PCB.

In conclusion, herein we propose an affordable and flexible wafer scale microfabrication process for hybrid multisite silicon/polyimide neural probes that can be adapted to multiple probe geometries and layouts according to in vivo experimental demands. The possibility of using interchangeable packaging with the same flexible interconnect cable design also permits great experimental flexibility and a reduction of fabrication and packaging costs. This way, we aim to contributing to a wider dissemination of neural probes and the empowerment of the neuroscience community to use sophisticated neuroengineering tools.

## Author Contributions

AN, CC and JG idealized the layout design and fabrication process. AN, HF and JF fabricated the Si probe. AN and LJ performed electrical impedance measurements and in vivo recordings. LJ analyzed the neural activity data. AN and LJ wrote the manuscript. JG and LJ financially supported this work. All authors have read and agreed to the published version of the manuscript.

## Funding

This work has been funded by National funds, through the Foundation for Science and Technology (FCT) - project UIDB/50026/2020 and UIDP/50026/2020; and by the projects NORTE-01-0145-FEDER-000013 (“PersonalizedNOS – New avenues for the development of personalized medical interventions for neurological, oncologic and surgical disorders”) and NORTE-01-0145-FEDER-000023 (“FROnTHERA – Frontiers of technology for theranostics of cancer, metabolic and neurodegenerative diseases”), supported by Norte Portugal Regional Operational Programme (NORTE 2020), under the PORTUGAL 2020 Partnership Agreement, through the European Regional Development Fund (ERDF).

## Acknowledgments

The authors would like to thank Dr. Patricia Monteiro, of the Life and Health Science Research Institute (ICVS), School of Medicine, University of Minho, for providing technical resources and for useful discussions.

## Conflict of interest

The authors declare no competing financial interests.

## Appendix A Neural probe structural characterization

SEM characterization was performed directly at wafer level without any preceding sample preparation, thus avoiding the introduction of artefacts into the measurements. SEM was performed to structurally evaluate the fabrication of the probes with a NovaNanoSEM 650 (FEI) imaging tool coupled with an Energy dispersive X-ray spectroscopy (EDX) Inca (Oxford instruments) detector.

To confirm if metal etch was successful in between metal lines, after metal patterning, SEM with EDX was performed (figure A1). Spectrum 1 taken from inter-line space shows that silicon (Si) and Aluminum (Al) are the most prevalent elements corresponding to the wafer composition and respective passivation. Other elements present at this step of the process, such as carbon (C) and oxygen (O), correspond to the photoresist used as mask for the metal dry etch and etching residues. Spectrum 2 was taken from the center of the electrode site and shows Au as the major element. Spectrum 3 was taken from a wall of a metal line which contains dry etch residues and photoresist mainly composed of C and O (figure A1 b and c) that posteriorly would be covered with the passivation layer.

**Figure A1.**
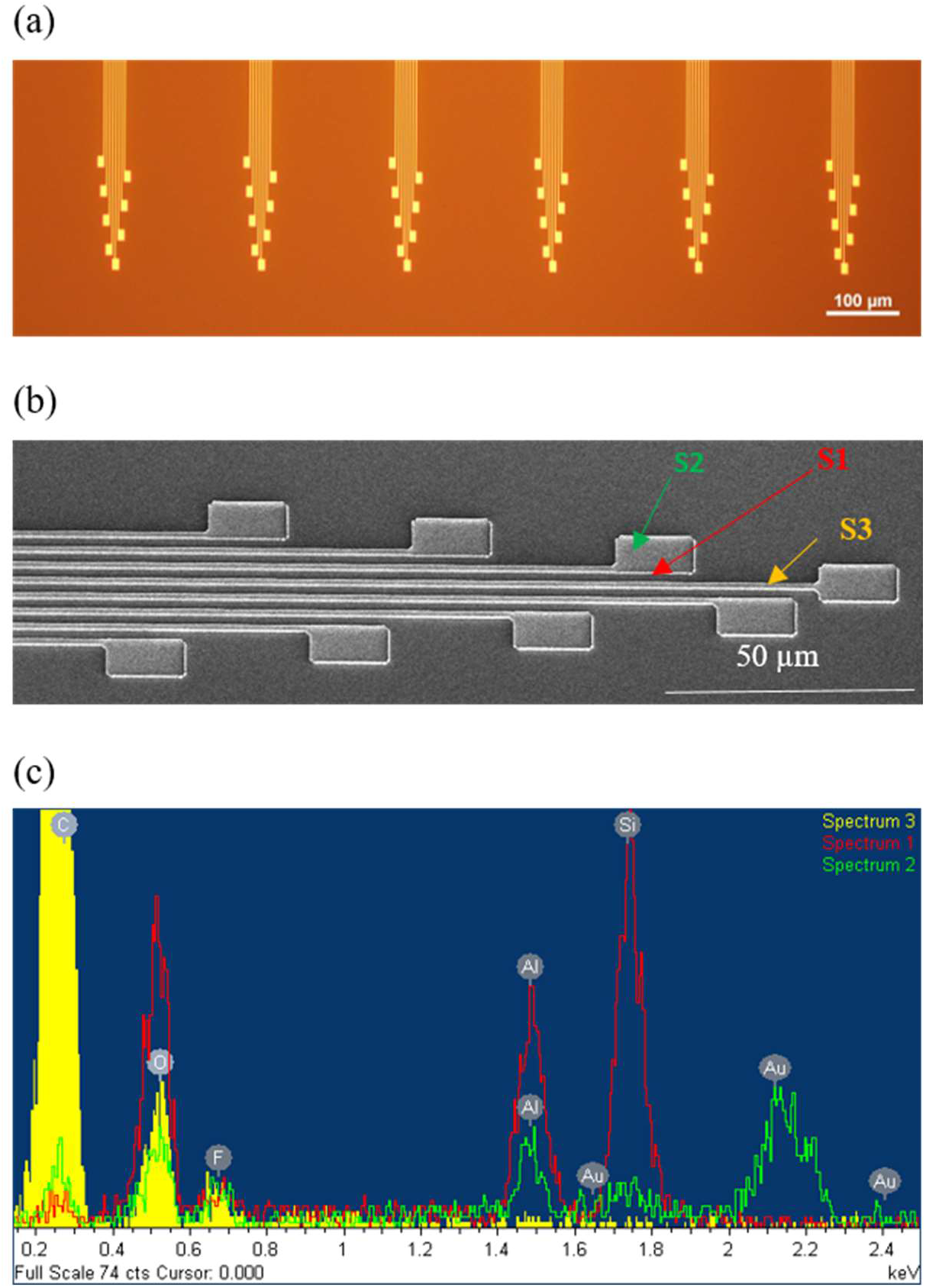
Structural characterization of electrode sites and interconnect lines: (**a**) Optical microscopic image of the electrode sites after metal patterning; (**b**) SEM image of the electrode sites after metal dry etch; (**c**) EDX to confirm metal etch in between interconnect metal lines (S1 – Si and Al_2_O_3_; S2 – Au; S3 – C).

## Notes

### Competing Interest Statement

The authors have declared no competing interest.

